# Mapping gene expression across the microbiome-gut-brain axis of germ- and specific pathogen-free mice

**DOI:** 10.1101/2025.04.15.648950

**Authors:** Clio Dritsa, Scott Hoffmann, Heather Hulme, Connor Lynch, Vicky Taylor, Richard Burchmore, John Cole, Richard J.A. Goodwin, Daniel M. Wall

## Abstract

Perturbation of the gut microbiota has been implicated in neurological diseases via communication across the microbiome-gut-brain axis. As a result, the discovery of mechanisms underlying interaction across this axis are becoming increasingly important. The germ-free (GF) mouse model has enabled an improved understanding of the influence of the gut microbiota on brain development and function. By utilising an advanced spatial profiling approach, we determined transcriptional changes in the brain, improving our understanding of how brain cells function and interact within their microenvironment in the absence of microbiome influence. Targeted regions of interest were selected based on brain regions implicated in neurological disease or reported structural differences between GF mouse brains and those of colonized mice. In the hippocampus 276 differentially expressed genes (DEGs) were identified, 345 DEGs in the thalamus, and 21 DEGs in pons. Contrastingly we identified only 2 DEGs in the midbrain and 4 in the medulla oblongata, with no DEGs in the cerebellum or corpus callosum. This data provides an overview of gut microbiota influence on gene expression in the brain, highlighting multiple genes of interest for further investigation in the context of microbiome influence on brain function and their potential relevance to neurological disease.

## Introduction

The gut microbiome is now known to exert significant effects on both mammalian physiology and the development of disease, while its modulation has been suggested to offer the promise of improving disease outcomes. Many central nervous system disorders, including neurodevelopmental disorders such as autism spectrum disorder, attention deficit hyperactivity disorder and neurodegenerative conditions such as Alzheimer’s and Parkinson’s disease, have had the gut microbiota implicated in their onset or pathology (1). However, while bi-directional communication mechanisms between the gut microbiota and the brain are mediated by many known host and microbial molecules, research on the microbiome-gut-brain (MGB) axis has failed to uncover mechanistic evidence linking the gut microbiota to induction or exacerbation of these aforementioned conditions.

Despite the limitations of studies using GF mice, they have significantly contributed to our understanding of microbiota-dependent effects on brain development, brain function and modulation of behaviour by the gut microbiota (2). Studies have demonstrated that GF animals exhibit increased blood brain barrier (BBB) permeability and defective neurogenesis (3, 4). Moreover, it has been shown that GF mice display distinct anatomical differences in the hippocampus with hypertrophic dendrites compared to colonised mice, hyper-myelinated axons in the prefrontal cortex, and both the abundance and maturity of microglia cells is altered alongside reduced myelination (5). GF mice also have altered stress responses and anxiety-like behaviour, disrupted metabolism and altered immune response as well as changes in neurotransmission (6). Increased anxiety and stress responses can be modulated by recolonisation of the intestine with probiotics, further underlining a role for the gut microbiota (7).

Understanding how the gut microbiota influences the brain is a significant challenge requiring novel tools and approaches. Our previous work has employed mass spectrometry imaging to determine the effects of the gut microbiota on neurotransmitter levels in GF and SPF mice (8, 9). Surprisingly, few changes were found in neurotransmitter levels in the gut and no significant changes were noted in the brains of GF mice in comparison to their colonised counterparts. However, changes in gene expression have been noted in GF mice across a number of studies, including through the use of single-cell RNA sequencing focused on the prefrontal cortex and hippocampus (10). The changes in microglial subtypes identified in GF mice mirror those in Alzheimer’s disease and major depressive disorder. Changes in the gut microbiota were also shown to influence expression of genes including those linked to neurodevelopment, stress response and cognition, such as emotional learning and memory, while the hippocampus and amygdala of GF mice displayed a changed expression profile of synaptic genes, leading to altered morphology, increased anxiety, and social impairments (5).

Therefore to complement our previous molecular mapping of the GF mouse brain here we undertook a spatial approach to understanding microbiota effects on neural transcription. Using a spatial transcriptomics approach across the GF murine brain, we identified regions of interest and directly compared them between GF and colonised specific pathogen free (SPF) mice. Our approach identified significant changes localized to specific regions of the brain, highlighting that certain brain regions are more susceptible to the absence of a gut microbiota.

## Materials and Methods

### Animal work

Six- to 8-week-old male C57BL/6J GF and SPF mice were sourced from the University of Manchester, Gnotobiotic Facility. Both GF and SPF mice were fed the same pelleted diet, which was sterilized by irradiation with 50 kGy. The Manchester Gnotobiotic Facility was established with the support of the Wellcome Trust (097820/Z/11/B) using founder mice obtained from the Clean Mouse Facility, University of Bern, Switzerland. The brain was removed from the skull and after removing the meninges, the brain was cut at the midline to separate the two hemispheres which were snap frozen in dry ice and isopropanol, followed by dry ice and isopentane for 30 seconds. Alcohol from tissues was allowed evaporate on dry ice prior to storing at -80°C. All work was carried out under UK Home Office license number PPL 7008584.

### Tissue processing

Brain sections were cut at -20°C at 10 μM. Multiple consecutive sagittal sections were thaw mounted on Superfrost glass slides, airdried using compressed air and vacuum sealed and stored at -80°C until further processing.

### GeoMx Spatial Transcriptomics

Slides were fixed overnight in 10% neutral buffer formalin, baked, and ethanol fixed to ensure tissues adhered to glass slides. For spatial transcriptomic profiling, we followed the semi-automated slide preparation method (Nanostring using a Bond RX (Leica Biosystems)). Slides were hybridised with the Mouse Whole Transcriptome Atlas (Nanostring, Cat#121401103) overnight at 37°C and subsequently stained with brain morphology markers and Styo13 (Table 1). Regions of interest (ROIs) were selected using the Nanostring DSP using various shapes, ensuring however that each ROI contained at least 200 cells (Supplementary Table 1). ROIs from the slides were collected into two separate 96-well plates and prepared for Illumina sequencing following the Nanostring library preparation protocol using the GeoMx Seq Code plates A+B (GeoMx Seq Code Pack A/B, Cat#121400201). Sequencing was performed on a NovaSeq6000 System (Illumina, San Diego, CA, USA).

**Table 1.** Fluorescent antibodies used for visualising the brain morphology and aiding the ROI selection.

| Target | Clone | Fluorophore |
| --- | --- | --- |
| Syto13 | Syto13 | Syto13 |
| GFAP | GA-5 | AF532 |
| MBP | Polyclonal | AF594 |
| NFH | NF01 | AF647 |

**Table 2.**
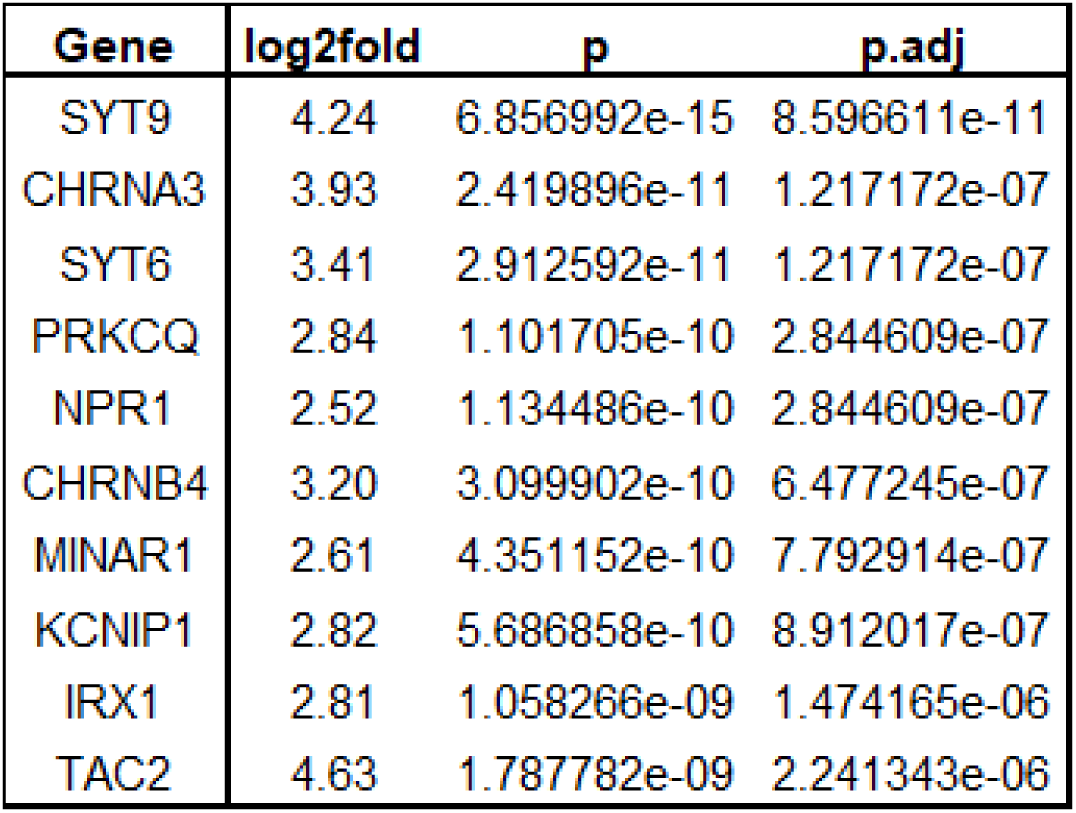
DEGs most significantly changed in GF compared to SPF hippocampus. Top 10 most significant DEGs in the GF compared to SPF hippocampus with cut off *p*. adj <0.05, absolute log2 fold > 0.5. All 10 DEGs were significantly increased in expression.

### Spatial Transcriptomic Data Analysis

The read count data was downloaded from the Nanostring platform, and downstream differential analysis performed in R. Firstly, samples that were technical replicates (same sample and ROI type) were merged by summing the read counts, and the surface areas. This provided a "weighted average" without unnecessarily removing observations (reads). The data was next corrected for surface area using SVA - Combat Seq (11). Next each pertinent pairwise differential comparison was performed using DESeq2 (12). The data was visualised using Searchlight specifying pairwise DE comparisons only, with significance at adjusted *p*< 0.05 (13). The Over Representation Analysis (ORA) function was used to identify enrichment of pathways, specifying the STRING 11.5 gene set database. The cut-off for pathway enrichment was adjusted *p* <0.05.

## Results

### ROI selection in SPF and GF mouse brain and PCA analysis of regional gene expression

Immunohistochemical staining of markers MBP, NF-H, GFAP and nucleic marker Syto13 was used to identify different brain structures and guide ROI selection for whole transcriptome profiling (Supplementary Figure 1). PCA analysis was initially used to observe the variation in gene expression between the different brain regions of the SPF brain to demonstrate that there is distinct gene expression in each brain region (Supplementary Figure 2).

### DEG analysis between SPF and GF brain structures

To identify DEGs between GF and SPF mice for different brain regions, pairwise comparisons were carried out with significance set at *p.* adj < 0.05. The number of significant DEGs was visualised using a lollipop chart indicating which genes increased or decreased in expression within each comparison (Figure 1). In the hippocampus we observed 276 DEGs, and 345 DEGs in the thalamus. Other regions had far fewer DEGS reflective of more limited influence of the microbiome. No significant DEGs were found for pairwise comparisons of GF versus SPF mice for the corpus callosum or cerebellum. Based on the number of significant DEGs, analysis focused on three ROIs with most gene expression changes, namely the hippocampus, the thalamus, and the pons. Our approach was to firstly carry out pairwise comparisons between GF and SPF mice for each brain region and then common, shared or unique DEGs were identified for these regions.

**Figure 1.**
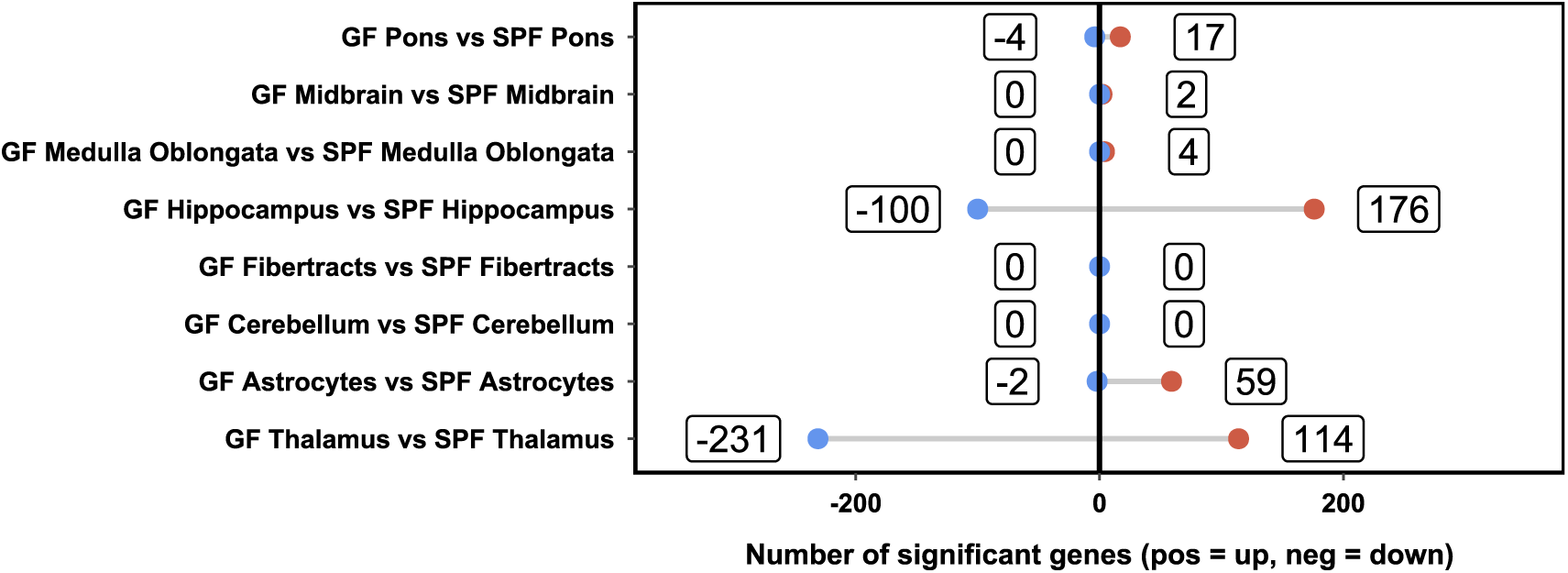
Altered gene expression in brain regions between GF and SPF mice. Lollipop plot showing the number of significant DEGs (*p*. adj <0.05, absolute log2 fold > 0.5) in pairwise differential expression comparison of GF and SPF mouse brain ROIs. Positive DEGs in red have higher expression in GF compared to SPF mice, negative DEGs shown in blue have lower expression in GF compared to SPF mice.

### Significant differences in gene expression between the GF and SPF hippocampi

To determine the effect of microbiota presence on hippocampal gene expression in mice, a differential expression analysis using DESeq2 was carried out. A volcano plot was used to visualize the relationship between fold change and the associated *p*- value for each gene. This plot validated a consistent relationship between the fold change and the *p*-value across all genes, as well as a balance between the direction of the changes between the two groups (Figure 2A). A hierarchically clustered heatmap was used to visualise and verify consistency among DEGs between replicates of each of the two groups. For this plot, genes were clustered by the similarity of their expression profiles and showed a pattern of expression among replicates (Figure 2B).

**Figure 2.**
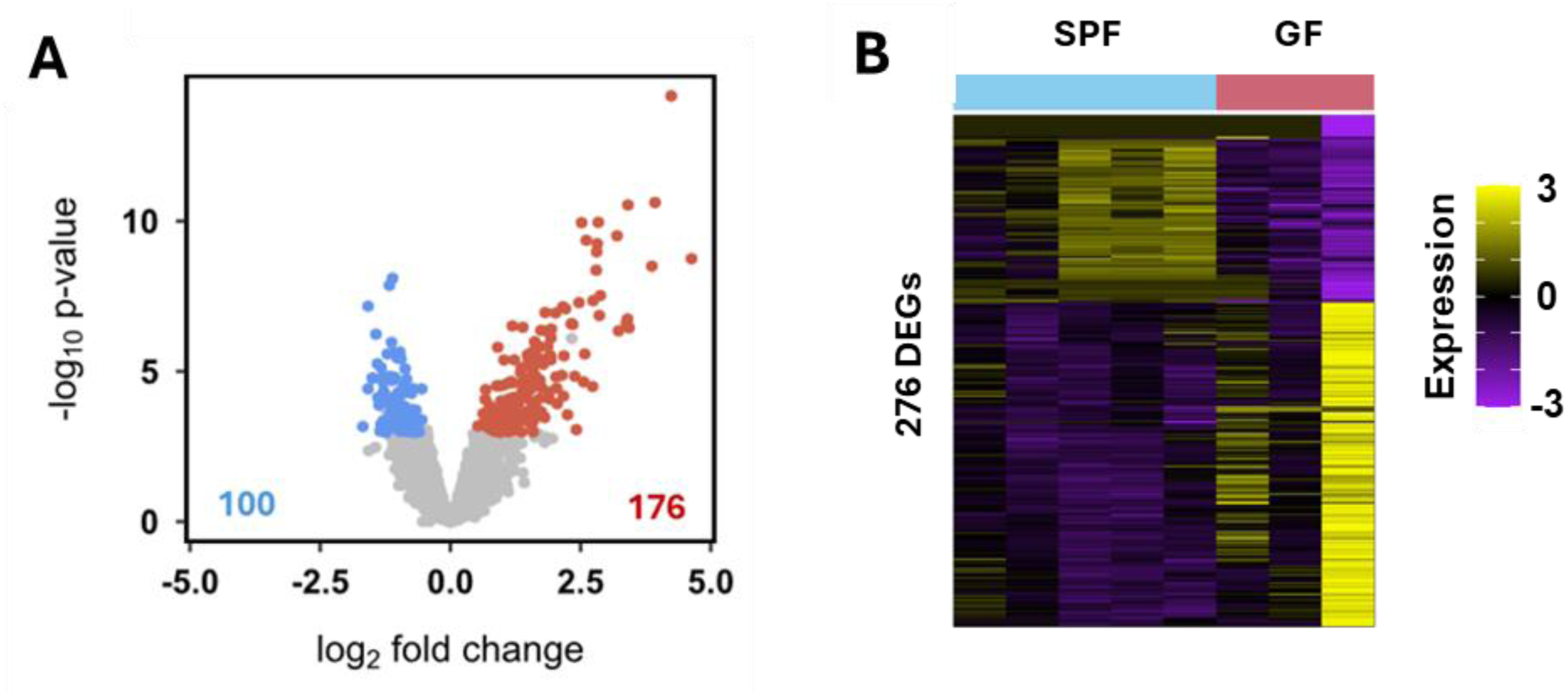
Identified DEGs between GF and SPF murine hippocampus. (**A**) Volcano plot illustrating significant DEGs (*p*. adj <0.05, absolute log2 fold > 0.5) for the comparison of the GF hippocampus and SPF hippocampus. (**B**) Hierarchically clustered heatmap of the significant DEGs between GF hippocampus and SPF hippocampus. Samples are on the x axis and DEGs on the y axis. Colour intensity represents expression level, with purple representing low expression, and yellow representing high expression. Expression levels have been row scaled into z−scores. The y-axis (both plots) and x-axis (right plot) have been hierarchically clustered using Spearman distances, with UPMGA agglomeration and mean reordering.

Following this, highly expressed DEGs in the GF hippocampus were compared to SPF hippocampus based on *p.* adj <0.05. This highlighted the most significantly altered expression profiles but revealed large log-fold changes in expression of genes, some of which were implicated in synaptic activity (Figure 3).

**Figure 3.**
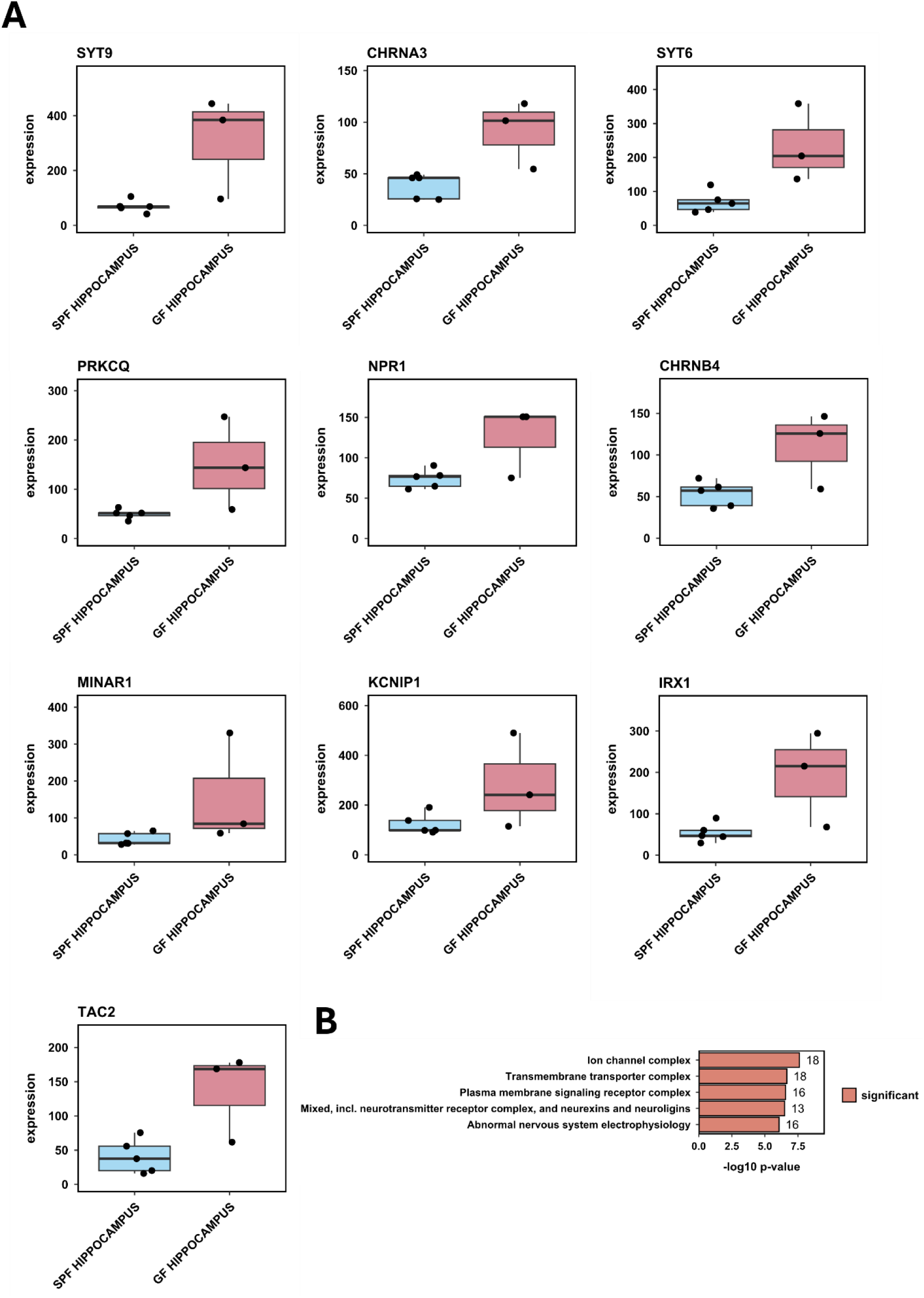
Boxplots of top ten significant DEGs in the GF compared to the SPF hippocampus and ORA for all significant hippocampal DEGs. (**A**) Boxplots showing GF hippocampus n=3 (red) and SPF hippocampus n=5 (blue) for the ten most significantly changed DEGs found in GF hippocampus (*p*. adj < 0.05, absolute log2fold > 0.5). Each dot is one sample, with sample groups shown on x-axis and gene expression values on the y-axis with mean (parallel line) and standard error (vertical to x-axis) for each sample group. (**B**) Summary bar chart of the 5 most enriched genes sets for all significantly differentially expressed genes (*p.* adj 0.05, absolute log2 fold > 0.5) between GF hippocampus and SPF hippocampus, significantly upregulated genes and significantly downregulated genes. For this plot the log10 enrichment *p*-value is on the x-axis and gene set name on the y-axis. Data labels represent the number of significant DEGs within each set. Enrichment analysis was performed using Hypergeometric Gene Set Enrichment on the gene set databases STRING11.5. Significantly enriched gene sets shown (*p*. adj < 0.05, absolute log2 fold > 0.0) are coloured in red.

Consideration was given to also examine DEGs that were expressed at lower levels in the GF compared to the SPF hippocampus based on *p*. adj <0.05. For this comparison it was observed that absolute log-fold changes were smaller than for highly expressed DEGs in the GF hippocampus, ranging from absolute log fold decrease of 0.97 to 1.58 (Supplementary Table 2). Reduced expression was seen in the GF hippocampus for genes such as *Gpr68*, a proton sensitive G protein coupled receptor, *Prox1,* which has been linked to adult neural fate specification, *Lrrtm4* involved in the regulation of synapse assembly, and *Cyp7b1* which is part of the cytochrome P450 family (Supplementary Figure 3).

### Gene enrichment analysis for DEGs between GF and SPF hippocampus of mice

A further aim of this analysis was to assess any global pathways or biological processes that the genes captured in the differential expression analysis may represent. To achieve this aim, an Over Representation Analysis (ORA) method was employed where the significantly DEGs were compared against a molecular signatures database. For this analysis STRING11.5 was used, which contains a priori defined list of gene sets corresponding to a distinct pattern such as a pathway, function or biological process. For this all significant DEGs between the GF hippocampus and SPF hippocampus were included in this analysis with *p*. adj <0.05 and absolute log2fold > 0.5 (Figure 3B). Using a hyper-geometric distribution test, the various gene sets comprising significant DEGs were assessed for frequency enrichment (more present or over representation) or depletion (under representation). The most enriched gene sets had the lowest *p*-values, as this confirmed that the gene set that was overrepresented (enriched) was not due to chance. This approach provided us with a useful snapshot of pathways or general biological concepts that are most affected by the DEGs and whether DEGs within each gene set are less or more expressed in each of the GF and SPF groups.

ORA analysis revealed functional categories that were enriched in the GF and SPF hippocampi, enriched categories included ion channel complexes and transmembrane transporter complexes (Figure 3B). Using these functional categories as guides, specific DEGs within these categories were further examined to determine their relative contribution to each enriched gene set (Supplementary Figure 4).

### Significant DEGs between the pons of GF and SPF mouse brains

The next comparison was between the GF pons and SPF pons, following the same analytical approach for the hippocampus sample analyses described above. To examine variance, a PCA plot was drawn but it showed little difference between the expression profiles of the two groups (Supplementary Figure 5). In addition, this showed more genes to have a positive log2 fold change (red dots), and a greater proportion of DEGs exhibited an increase in the GF samples (Figure 4B). Overall, there were 21 DEGs identified when comparing GF and SPF pons, with 17 being highly expressed in GF mouse pons and four being lower expressed in the pons of GF mice (Figure 4A). The heatmap displays the pattern of gene expression within each of the biological samples in this study (Figure 4B).

**Figure 4.**
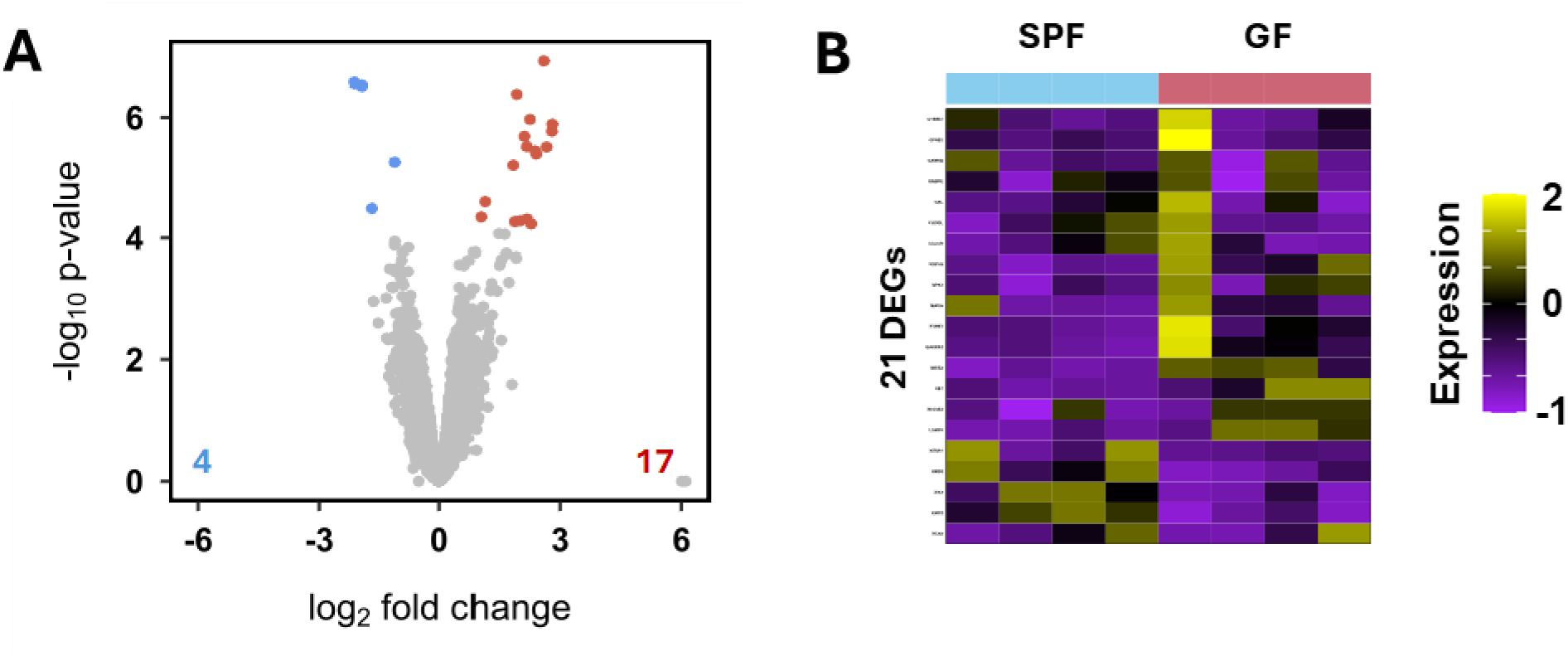
Identified DEGs between the GF and SPF murine pons. (**A**) Volcano plot illustrating significant differential genes (*p.* adj <0.05, absolute log2 fold > 0.5 for the comparison of GF pons vs SPF pons. (**B**) Hierarchically clustered heatmap of the significantly differentially expressed genes (*p.* adj < 0.05, absolute log2 fold > 0.5) between GF pons and SPF pons. Samples are on the x-axis and genes on the y-axis. Colour intensity represents expression level, with purple representing low expression, and yellow representing high expression. Expression levels have been row scaled into z−scores. The y-axis (both plots) and x-axis (right plot) have been hierarchically clustered using Spearman distances with UPMGA agglomeration and mean reordering.

We explored DEGs between the GF and SPF mouse pons based on *p.* adj <0.05 and absolute log2 fold (Figure 5A; Table 3). Table 3 shows *p*-value and absolute log2 fold change larger than 1.9-fold for the top ten DEGs between GF pons and SPF pons. Following this, we were interested in the DEGs that had significantly decreased in GF pons compared to SPF pons. Decreased genes had not changed to the same degree as those that were found to be increased with only 2 genes showing a corresponding 1.9-fold or greater decrease (Supplementary Table 3; Supplementary Figure 6). One such gene with significantly lower expression in GF pons when compared with SPF pons was the *Six3* gene (1.9-fold), implicated in signalling pathways related to retinal development and required for the development of the forebrain (14).

**Figure 5.**
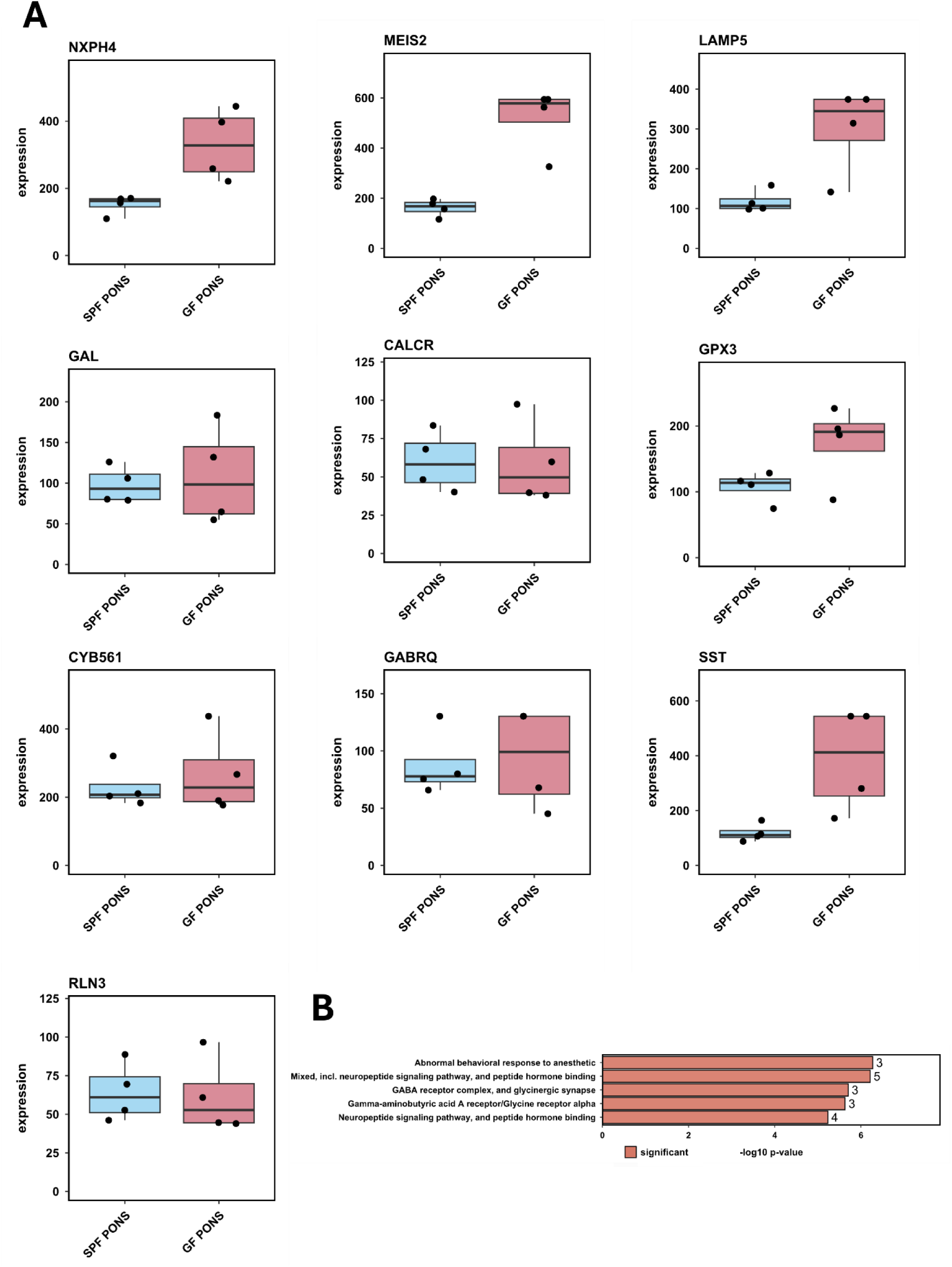
Boxplots of top ten DEGs with significantly higher expression in the GF compared to the SPF pons and ORA of all significant DEGs indicating enriched functional categories. **(A)** Boxplots of gene expression values of GF pons n=4 (red) and SPF pons n=4 (blue) for the 10 most significantly changed DEGs found in GF pons, (*p*. adj < 0.05, absolute log2fold > 0.5). Each dot is one sample, with sample groups shown on x-axis and gene expression values on the y-axis with mean (parallel line) and standard error (vertical to x-axis) for each sample group. (**B**) Summary bar chart of the top five most enriched genes sets for all significantly differentially expressed genes (*p.* adj 0.05, absolute log2 fold > 0.5) between GF pons and SPF pons, significantly upregulated genes and significantly downregulated genes. For this plot–log10 enrichment *p*-value is on the x-axis and gene set name on the y-axis. Data labels represent the number of significantly differential expressed genes within each set. Enrichment analysis was performed using Hypergeometric Gene Set Enrichment on the gene set databases STRING11.5. Significantly enriched gene sets (*p.* adj < 0.05, absolute log2 fold > 0.0) are coloured red.

**Table 3.** Top 10 DEGs significantly increased in expression in the GF compared to SPF pons. Ten most significantly increased DEGs with cut off *p*. adj <0.05, absolute log2 fold >0.5.

| Gene | log2fold | p | p.adj |
| --- | --- | --- | --- |
| NXPH4 | 2.59 | 1.206315e-07 | 0.001446376 |
| MEIS2 | 1.91 | 4.116134e-07 | 0.001467813 |
| LAMP5 | 2.24 | 1.073878e-06 | 0.003063558 |
| GAL | 2.80 | 1.290951e-06 | 0.003069021 |
| CALCR | 2.79 | 1.685559e-06 | 0.003434687 |
| GPX3 | 2.10 | 2.053106e-06 | 0.003660687 |
| CYB561 | 2.16 | 3.019228e-06 | 0.004389102 |
| GABRQ | 2.66 | 3.077049e-06 | 0.004389102 |
| SST | 2.37 | 3.579025e-06 | 0.004641019 |
| RLN3 | 2.40 | 4.018089e-06 | 0.004776168 |

### Gene enrichment analysis among all significant DEGs between the GF pons and SPF pons

Using STRING gene-set data to obtain gene ontology terms, ORA was performed on significant DEGs between GF pons and SPF pons. The functional classifications revealed were related to abnormal behavioural response to anaesthetic, neuropeptide signalling and peptide hormone binding, GABA receptor and glycinergic synapse (Figure 5B).

### Significant DEGs between the GF thalamus and SPF thalamus

The final pairwise comparison was between the GF thalamus and SPF thalamus, following the same steps as for the analysis of the hippocampus and pons above. The PCA plots showed variance between samples with distinct expression patterns between GF thalamus and SPF thalamus as the two groups clustered separately (Supplementary Figure 7). The volcano plot displayed 231 DEGs to have a negative log2 fold change (blue dots) and 114 DEGs with a positive log2 fold change (red dots), showing that a greater proportion of DEGs exhibited a decrease in the GF thalamus when compared to the SPF thalamus (Figure 6A). A heatmap demonstrated the pattern of expression of the genes within each of the biological samples in this study (Figure 6B).

**Figure 6.**
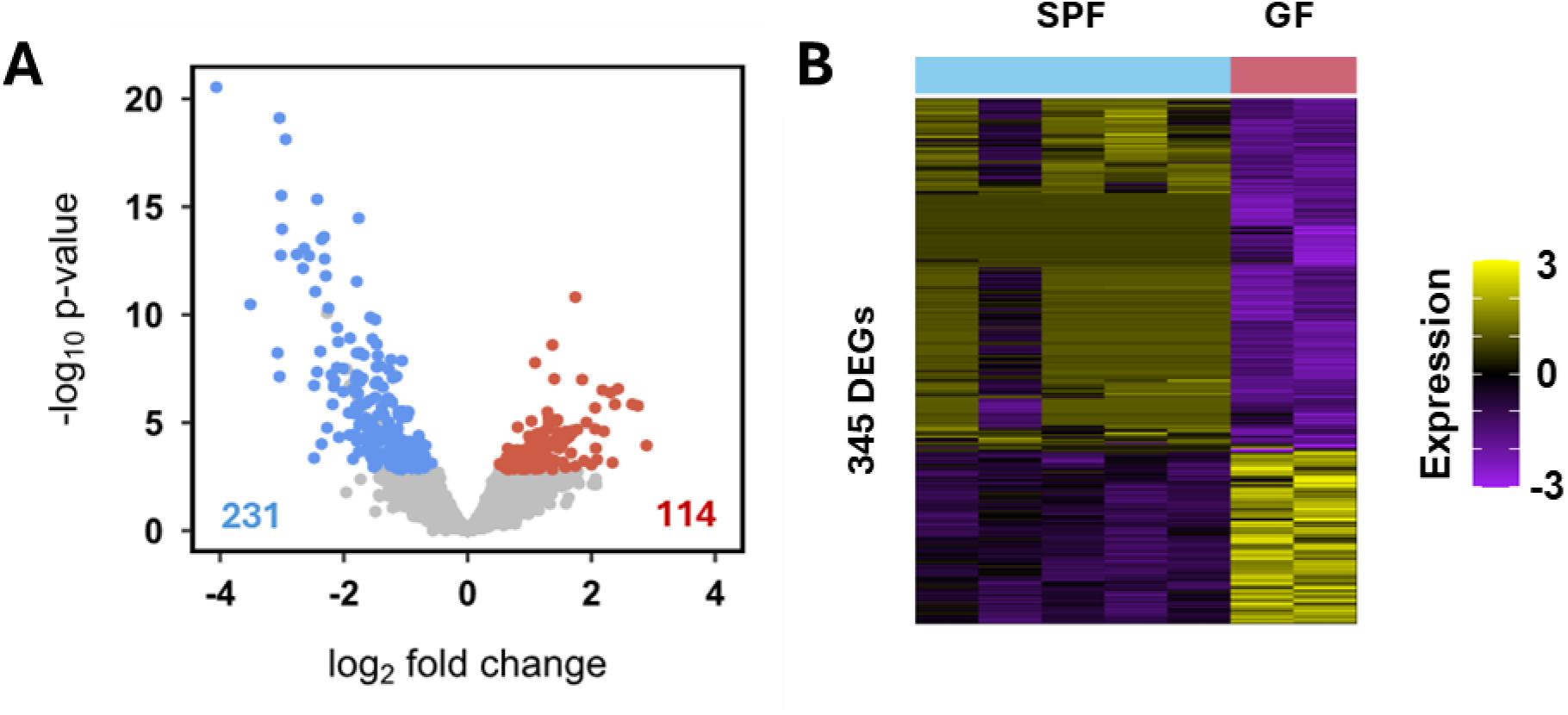
Significant DEGs between the GF and SPF murine thalamus. (**A**) Volcano plot illustrating significant DEGs (*p.* adj <0.05, absolute log2 fold > 0.5 for the comparison of GF thalamus versus SPF thalamus. (**B**) Hierarchically clustered heatmap of the significantly DEGs (*p.* adj < 0.05, absolute log2 fold > 0.5) between GF thalamus and SPF thalamus. Samples are on the x-axis and genes on the y-axis. Colour intensity represents expression level, with purple representing low expression and yellow representing high expression. Expression levels have been row scaled into z−scores. The y-axis (both plots) and x-axis (right plot) have been hierarchically clustered using Spearman distances with UPMGA agglomeration and mean reordering.

Furthermore, we looked at DEGs from the GF thalamus compared to the SPF thalamus (Figure 7) based on *p.* adj <0.05 and absolute log2 fold. Table 4 shows *p*- value and absolute log2 fold changes ranging from 1.09 to 2.66 for the top ten DEGs between the GF thalamus and SPF thalamus. DEGs in the comparison of the GF thalamus and SPF thalamus included the *Traf3* gene which displays a higher expression in the GF thalamus (Figure 7). This gene encodes an immune regulator found with the retina and its neuronal expression is thought to be linked with ischemic damage (15). We were also interested in the DEGs with a lower expression in GF thalamus compared to SPF thalamus (Supplementary Figure 8). *Gabra4* showed a significantly lower expression in the GF thalamus when compared to the SPF thalamus (Supplementary Table 4). *Gabra4* encodes for GABA receptor A4, the function of which is implicated in neurotransmission and changes within this system are associated to neurological conditions (16).

**Figure 7.**
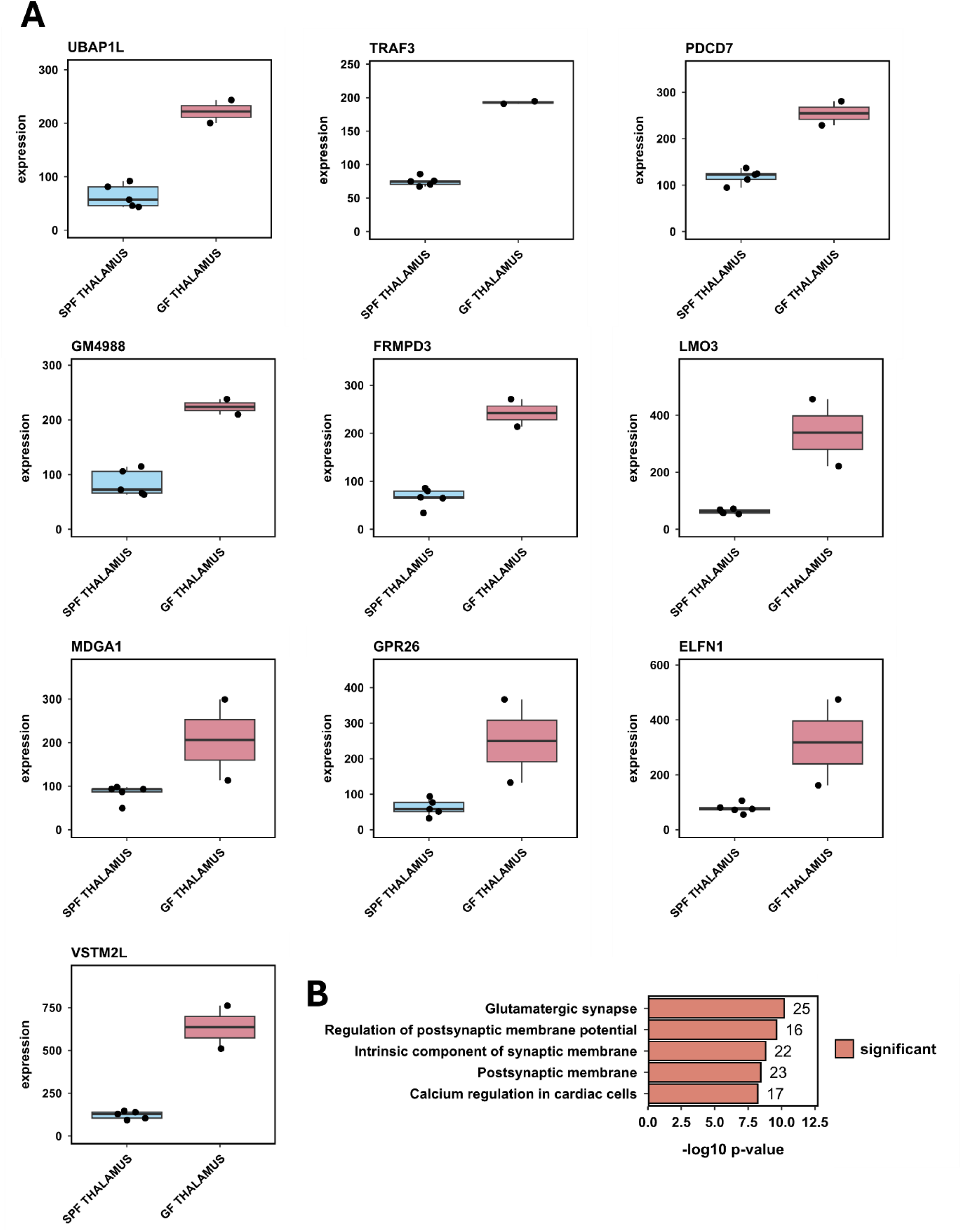
Boxplots of top ten DEGs with significantly higher expression in the GF compared to the SPF thalamus and ORA of all significant DEGs between GF and SPF thalamus indicating enriched functional categories. (**A**) Boxplots of gene expression value of GF thalamus n=2 (red) and SPF thalamus n=5 (blue) for the ten most significantly changed DEGs found in GF thalamus by adjusted *p*-value, (*p*< 0.05). Each dot is one sample, with sample groups shown on x-axis and gene expression values on the y-axis with mean (parallel line) and standard error (vertical to x-axis) for each sample group. (**B**) Summary bar chart of the top five most enriched genes sets for all significantly DEGs (*p.* adj 0.05, absolute log2 fold > 0.5) between the GF thalamus and the SPF thalamus, significantly upregulated genes and significantly downregulated genes. For this plot–log10 enrichment *p*-value is on the x-axis and gene set name on the y-axis. Data labels represent the number of significant DEGs within each set. Enrichment analysis was performed using Hypergeometric Gene Set Enrichment on the gene set databases STRING11.5. Significantly enriched gene sets (*p.* adj < 0.05, absolute log2 fold > 0.0) are coloured red.

**Table 4.** Top 10 DEGs significantly increased in expression in the GF thalamus compared to the SPF thalamus. Most significant DEGs with cut off *p*. adj <0.05, absolute log2 fold >0.5.

| Gene | log2fold | p | p.adj |
| --- | --- | --- | --- |
| UBAP1L | 1.74 | 1.536371e-11 | 9.439787e-09 |
| TRAF3 | 1.37 | 2.510309e-09 | 1.010529e-06 |
| PDCD7 | 1.09 | 1.665759e-08 | 5.117389e-06 |
| GM4988 | 1.40 | 9.432659e-08 | 2.079361e-05 |
| FRMPD3 | 1.85 | 1.010838e-07 | 2.185282e-05 |
| LMO3 | 2.43 | 2.717660e-07 | 5.035867e-05 |
| MDGA1 | 2.18 | 3.123886e-07 | 5.610500e-05 |
| GPR26 | 2.31 | 4.130340e-07 | 7.090822e-05 |
| ELFN1 | 2.66 | 1.387998e-06 | 2.038952e-04 |
| VSTM2L | 2.38 | 1.423083e-06 | 2.050996e-04 |

### Gene enrichment analysis among all DEGs between the GF thalamus and the SPF thalamus

ORA analysis was performed on all the DEGs from the pairwise analysis between the GF thalamus and SPF thalamus (Figure 7B). The top five most enriched biological functional categories identified included the glutamatergic synapse, the postsynaptic membrane and regulation of postsynaptic membrane potential, intrinsic components of the synaptic membrane, and calcium regulation of cardiac cells pathway.

### Determination of DEGs common to the pons, the hippocampus and the thalamus

Given their higher number of DEGs three brain regions were focused on for further analysis, the pons, the hippocampus and the thalamus. Common or unique DEGs among the three structures were then identified and a Venn diagram created to showcase any overlap among DEGs between these brain regions (Figure 8). There were large differences in the number of unique DEGs identified in the pons (20 DEGs) compared to the thalamus (320 DEGs) or hippocampus (252 DEGs) when comparing GF and SPF mice. No common DEGs were identified between the pons, the hippocampus and the thalamus, however, a single DEG was shared between the pons and the thalamus, *Ptprt,* a member of the protein tyrosine phosphatase family which is important for signalling in a multitude of cellular processes. A total of 24 DEGs were found to be common between the thalamus and the hippocampus. These DEGs are implicated in neuronal activities with, for example, a known association between *Prkcd* and G*ml1* in hippocampal memory signalling (17).

**Figure 8.**
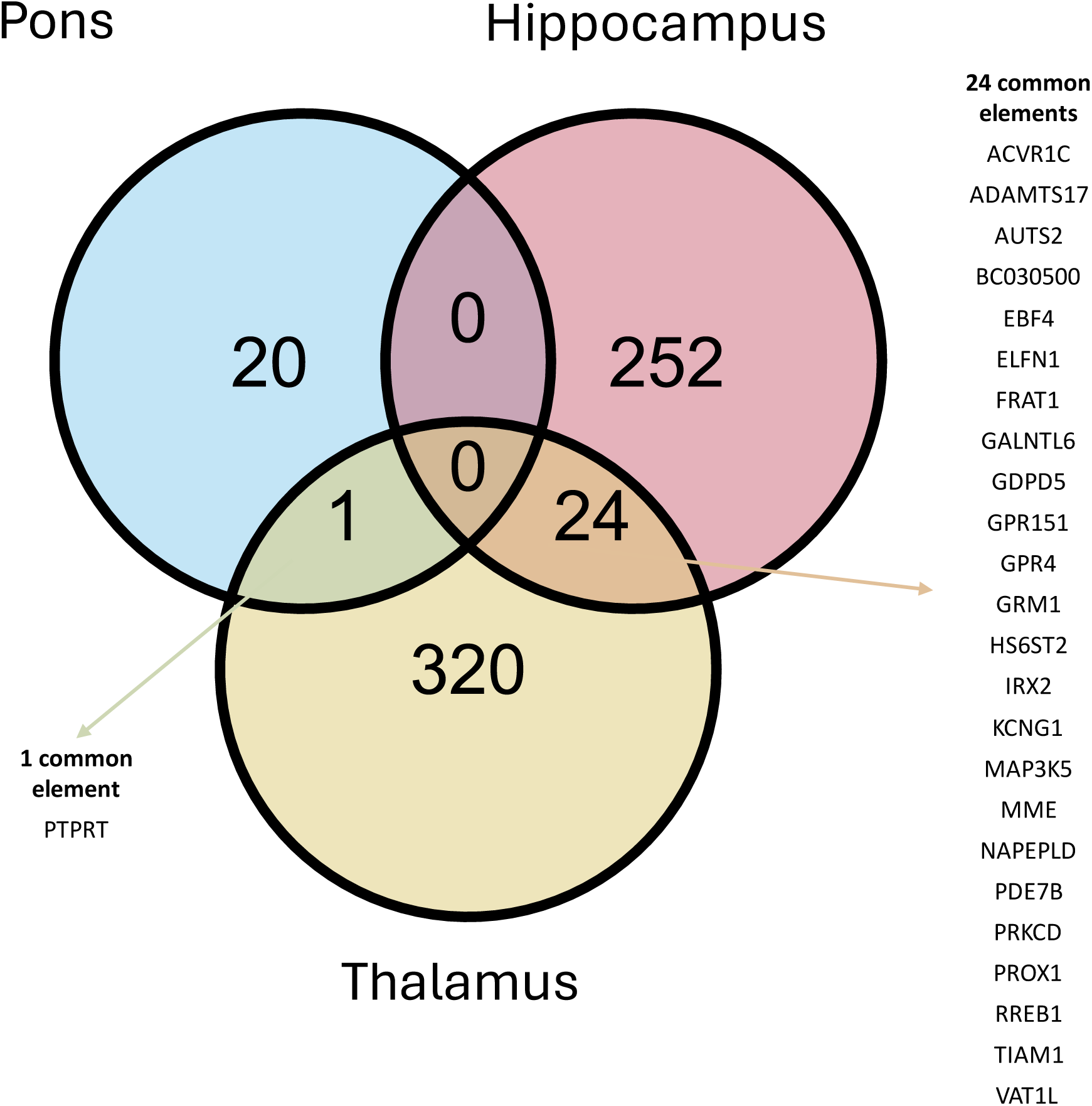
Venn diagram of three sets of significant DEGs between the pons, the thalamus and the hippocampus in GF and SPF mice. Venn diagram to capture any common genes shared between the pons, the thalamus and the hippocampus. No common DEGs were shared among all three ROIs. Overlapping significant DEGs are listed for the comparison of pons and thalamus (1 element), *p* <0.01 and for the comparison of hippocampus and thalamus (24 elements), *p* <0.01.

## Discussion

Studies conducted on GF animals have advanced our understanding of the dynamic interplay between the absence of a microbiota and quantifiable phenotypic changes, including molecular alternations (2, 3). Multiple aspects of nervous system physiology are altered in GF mice, which exhibit phenotypic differences in comparison to conventional mice, namely in blood-brain-barrier permeability, neurogenesis patterns, hypothalamic-pituitary-axis, stress response, and anxiety (2, 3, 18, 19). However studies using GF mice alongside a growing body of research on animal behaviour has also contributed to our understanding the MGB axis (6).

Evidence points to distinct microbiota profiles that are associated with cognitive function and behaviour, and how a lack of microbiota not only is correlated with impaired cognition, but also disease outcomes for depression and anxiety related behaviours (19). Additionally, an abundance of microbial genera in the gut have been associated with behavioural attributes in humans, where lower diversity was linked to higher stress levels and anxiety, with these characteristics linked to interindividual compositional differences in the gut microbiome (20).

To interrogate communication across the MGB-axis we applied spatial transcriptomics to generate a profile of gene expression in sagittal sections of mouse brain from SPF and GF mice. Brain transcriptome changes between SPF and GF showed numerous genes that were significantly changed per region which associated with significantly altered pathways. The differing number of DEGs seen in the brain regions selected, highlight how distinct brain regions are influenced differently by the gut microbiota. Previous research has highlighted regulation of hippocampal processes via MGB-axis signalling, with hippocampal neurogenesis, which is important for learning and memory and mitigation of stress responses in adulthood, modulated via the microbiota in adult mice, alongside morphological alterations in dendrites of neurons and transcriptional changes (21, 22).

Here we focus on some example genes identified in the different brain regions where the most significant number of DEGs were identified, namely the hippocampus, thalamus and pons. *Gpr68* a proton-sensing receptor that stimulates multiple cellular activities, was significantly decreased in the GF mouse hippocampus. In the hippocampus, *Gpr68* is widely expressed in particular within cornu ammonis (CA) 3 pyramidal neuronal cells (23). Using a *Gpr68* knockout mouse model, a reduction of fiber volley (FV) was demonstrated *ex vivo* upon stimulation of the Schaffer collateral-CA1 synapse in the hippocampus and its reduction was thought to contribute to deficits in avoidance memory based on negative reinforcement (foot shock) in an *in vivo* passive avoidance test (23). It was additionally demonstrated that *Gpr68* contributes to synaptic transmission given the reduced long-term potentiation measurements in the *Gpr68* knockout model.

Previous research suggested that the stimulation of the amygdala may lead to the regulation of synaptic plasticity in the hippocampus (24). Interestingly, protons and acid-sensing ion channels are thought to be essential components in synaptic plasticity in amygdala-related learning and memory as demonstrated by regulation of amygdala long-term potentiation (25). Changes in extracellular protons during synaptic transmission have been shown to inhibit N-methyl-D-aspartate (NMDA) receptors (primary excitatory neurotransmitter receptors) in cerebellar neurons (26). Studies on GF mice have also shown a decrease in the expression of NMDA receptor NR2B subunit in the cortex and hippocampus (19). Considering *Gpr68* is an acid ion receptor sensitive to extracellular pH changes, and is an integral component in synaptic transmission function, the *Gpr68* under-expression seen in our study could pose an obstacle in promoting optimal hippocampal function in GF mice.

The gut microbiota has been shown to influence stress responsiveness and modulate hypothalamic-pituitary-adrenal axis activity in GF mice (6, 19). GF mice display lower anxiety behaviour versus conventionally colonised mice, however they do exhibit higher levels of anxiety-like behaviour when levels of adrenocorticotropic hormone (ACTH) and cortisol are increased, which is reversible in young mice upon microbial colonisation (2). This highlights the concept of a critical developmental time window early in life, where neuronal plasticity could be influenced via the gut microbiota.

*Tac2* was found to be significantly increased in the GF hippocampus. *Tac2* is a member of the tachykinin family of neuropeptides, with *Tac1* encoding substance P and neurokinin A, whereas *Tac2* encodes for neurokinin B (NkB) (27, 28). Tachykinin was found to be widely expressed in neurons in the CNS, particularly in the emotional and social behaviour centres of the brain, namely the amygdala and hypothalamus. *Tac2* is also associated with several genes that play a part in hippocampal stress signalling and neural plasticity and consequently is also thought to be involved in these cognition-related processes (29). Studying *Tac2* expression and interaction with other genes in a gene networking investigation, *Tac2* expression was found to be significantly elevated in mice following prolonged exposure to mild stress conditions. In the same study, pathway enrichment analysis revealed a strong relationship between *Tac2* and neuroactive ligand-receptor interaction pathways such as those connected to the process of neural plasticity and genes within the calcium signalling pathway (29). In chronic social isolation, stress was shown to cause several behavioural changes in mice which were paired with overexpression of *Tac2* in multiple brain regions including the hippocampus (30). Further studies on *Tac2* and its receptor in the amygdala showed these were essential in altering fear memory consolidation and were also highly expressed after fear conditioning tests (31). This work highlights that tachykinin is stress-sensitive and necessary for diverse hippocampal processes such as cognition and fear memory consolidation. When examining the pons, there was a small number of DEGs in the GF versus SPF mice including genes relevant to brain development, such as the homeobox *Six3* gene, a key transcription factor involved in activation and suppression of genes (32). *Six3* is also regulated by SOX2, a transcription factor pivotal to maintaining self-renewal and pluripotency capacity of undifferentiated embryonic stem cells (33, 34). In the thalamus we identified a multitude of DEGs to be altered between the GF and SPF mice. Of interest was the GABA receptor A4 (*Gabra4*) receptor, which had decreased expression levels in the GF thalamus. GABA is the main inhibitory neurotransmitter of the central nervous system, mediating its effect through receptors specific to it including ionotropic GABA_A_ and GABA_B_ receptors, both of which are essential pharmacological targets. Changes within the GABAergic system have been implicated in psychiatric conditions (16). This was particularly noteworthy as this system has previously been documented to be altered in neurodevelopmental disorders (35). This finding could potentially further reinforce the role of the microbiome in modulating host gene expression in the context of neurodevelopmental disorders (36).

The overlap of the identified DEGs across the three brain regions focused on was also investigated. This analysis did not show a common DEG for all three regions, but a shared DEG between the pons and the thalamus and 24 overlapping DEGs between the thalamus and the hippocampus. The *Ptprt* gene, a member of the protein tyrosine phosphatase family, was found to be common between the thalamus and pons DEGs. The tyrosine phosphatase family is important signalling molecules in a multitude of cellular processes and *Ptprt* is linked signal transduction and cell adhesion in particular (37). PTPRT is thought to be vital for synaptic neurotransmission and neuronal dendritic spine formation by interaction with cell adhesion molecules and dephosphorylation of synaptic components (38). However, considering the role of the thalamus and the pons in coordination and support of other key brain functions, and the microbiomes association to neurological disease, *Ptprt* gene expression changes at the synapse level in our study may impact on synaptic plasticity and therefore contribute to altered neuronal activity.

We also identified gene signatures comprised of DEGs found in GF and SPF hippocampus and thalamus ROIs. This analysis highlighted enriched pathways such as gated channel activity and synaptic transmission, among others, which was not surprising considering previous reports on altered neurotransmission in GF mice and expression changes of synaptic genes. DEGs such as *Tac2* in the hippocampus, *and Gabra4* in the thalamus, are all implicated in neurotransmission. Also, the aforementioned enriched pathways were captured in over representation analysis of individual brain regions. Motor activity has been found to be increased in GF compared to SPF mice whereas behavioural differences were also seen with GF mice displaying a reduced anxiety phenotype (6). These findings strengthen existing links describing the influence of the gut microbiota on the brain gene transcription (2, 5, 10, 39–41).

Future studies will be refined by focusing on finer substructures within regions such as the hippocampus, as it would be interesting to interrogate gene expression changes within specific areas of the hippocampus and also globally within the hippocampus. Considering the amygdala is the emotional processing centre within the brain and a multitude of behavioural studies have focused on this structure, it would be useful to investigate if gene expression changes in the amygdala could be associated to behavioural traits previously reported in the context of an absent microbiome. Whilst further research is required to confirm the microbiome’s role in these changes, there appears to be an effect on gene expression between GF and SPF mice. These data also allow us to appreciate the dynamic, multifaceted interactions between the gut, microbiome, and brain gene expression, and potentially highlight target genes that may be useful in the context of neurological disease interventions.

## Supporting information

Dritsa et al. Supplementary Materials

