## Supplementary material for "Mapping gene expression across the microbiome-gut-brain axis of germ- and specific pathogen-free mice": Dritsa et al. Supplementary Materials

**Supplementary Figure 1**

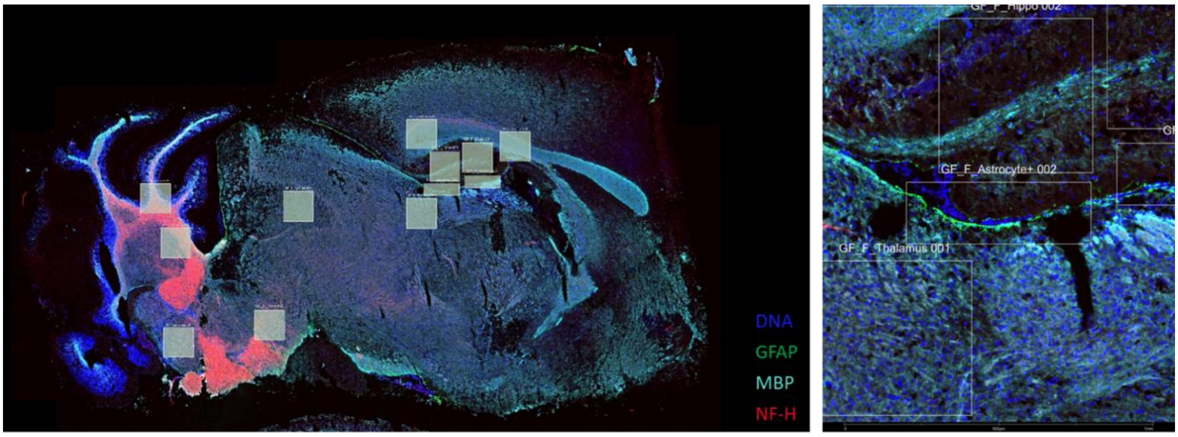


**Supplementary Figure 1. Multiplex Immunofluorescence (mIF) image taken on the Digital Spatial Profiler (DSP) platform.** Representative image of a 10 µM sagittal section of adult mouse brain using GFAP (green), MBP (light blue), NFH (red) and nucleic acid immunofluorescence marker (dark blue) captured on the Nanostring DSP for morphological identification of structures. The mask square boxes indicate ROIs that were profiled on the Nanostring GeoMx platform using the GeoMx Mouse Whole Transcriptome Atlas.

#### Supplementary Figure 2

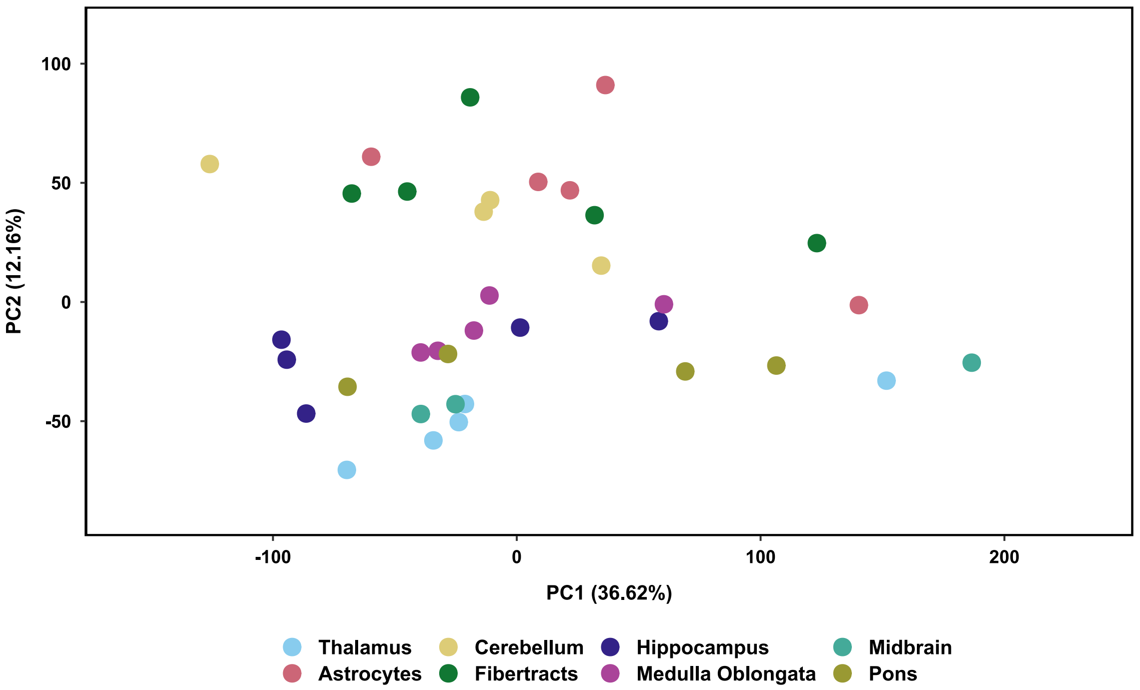


**Supplementary Figure 2. Assessing variation within the data by PCA plot.** PCA of gene expression in specific brain regions of a representative SPF mouse brain.

#### Supplementary Figure 3

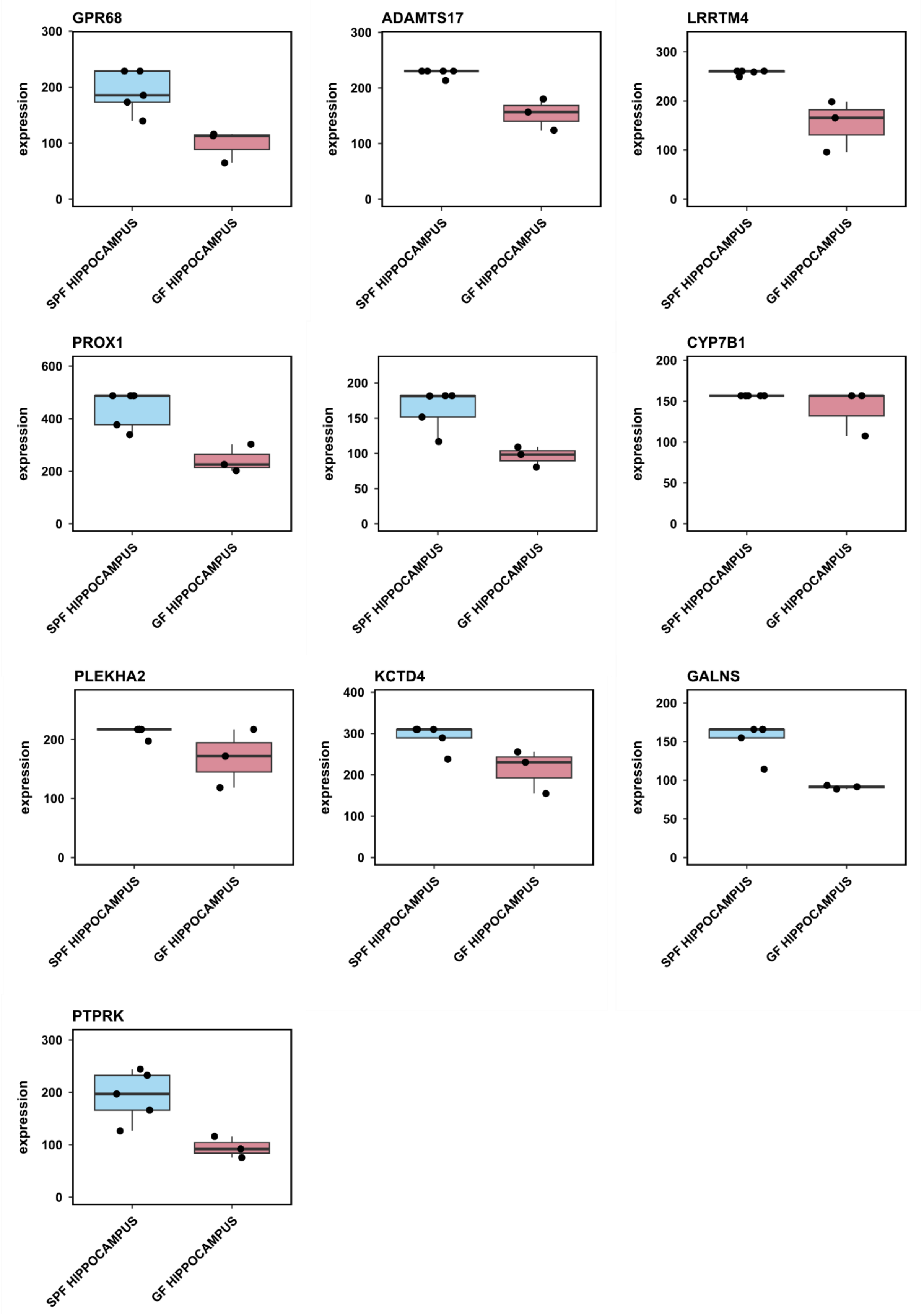

**Supplementary Figure 3. Boxplots of top ten DEGs with significantly lower expression in the GF compared to the SPF hippocampus.** Boxplots showing GF hippocampus n=3 (red) and SPF hippocampus n=5 (blue) for the ten most significantly decreased DEGs in GF hippocampus (*p*. adj < 0.05, absolute log2fold > 0.5). Each dot is one sample, with sample groups shown on x-axis and gene expression values on the y-axis with mean (parallel line) and standard error (vertical to x-axis) for each sample group.

**Supplementary Figure 4**

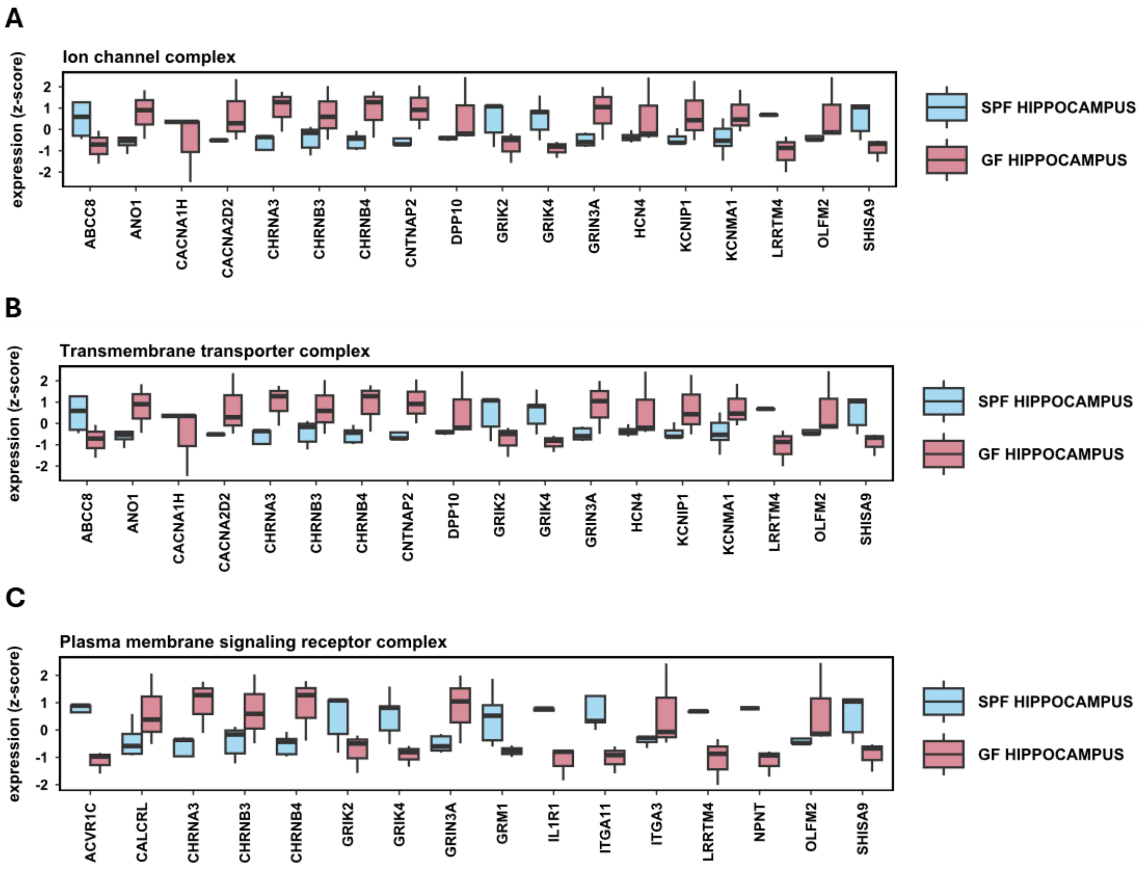


**Supplementary Figure 4. ORA summary boxplots of DEGs for the top three most enriched gene sets for the comparison of GF and SPF hippocampus.** Representative summary boxplots with expression values in z-scores to illustrate the three top, most enriched genes sets for all significantly DEGs (*p.* adj 0.05, absolute log2 fold > 0.5) between GF hippocampus and SPF hippocampus, significantly upregulated genes and significantly downregulated genes. (**A**) Ion channel complex gene set. (**B**) Transmembrane transporter complex gene set. (**C**) Plasma membrane signalling receptor complex gene set. For each plot the significantly differential genes within each gene set are provided on the x-axis and expression values scaled into per gen z-scores are shown on the y-axis. Boxes are shown for GF hippocampus (red) and SPF hippocampus (blue) groups. Enrichment analysis was performed using Hypergeometric Gene Set Enrichment on the gene set databases STRING11.5. The figure provides a visual representation of the DEGs comprising 3 out of 5 gene sets taken from Figure 3B, namely the ion channel complex, the transmembrane transporter complex and the plasma membrane signalling receptor complex. It was observed that the DEGs within these gene-sets were primarily highly expressed in the GF hippocampus versus SPF hippocampus. The DEGs between ion channel complexes and transmembrane transporter complexes gene sets were shared (seen in Figure 8A and Figure 8B). The plasma membrane signalling receptor gene set (Figure 8 C), comprised many of the DEGs seen in the first and second most enriched categories (Figure 8A and Figure 8B) such as *Chra3*, *Chrnb3*, *Chrnb4*, *Grik2*, *Grik4*, *Grin3a*, *Llrtm4*, *Olfm2* and *Shisa9*. However, the plasma membrane signalling receptor gene set also comprised DEGs that were not seen in the first two enriched categories, namely *Acvr1c*, *Calcrl*, *Grm1*, *IL1R1*, *Itga11*, *Itga3* and *Npnt*. While individual DEGs showed higher log fold changes (Table 2), ORA analysis indicated that DEGs within functional categories of genes sets were significantly changing, but at lower levels (Figure 8).

**Supplementary Figure 5**

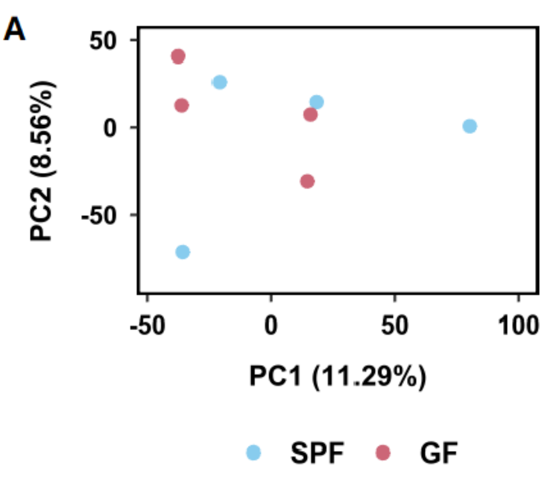


**Supplementary Figure 5. Clustering of DEGs from comparison of the GF and SPF pons.** PCA plot of GF pons n=4 (red) and SPF pons n=4 (blue), showing the first principal component (PC1) accounting for 50.44 % of variance and the second principal component (PC2) capturing 20.87% of variance.

#### Supplementary Figure 6

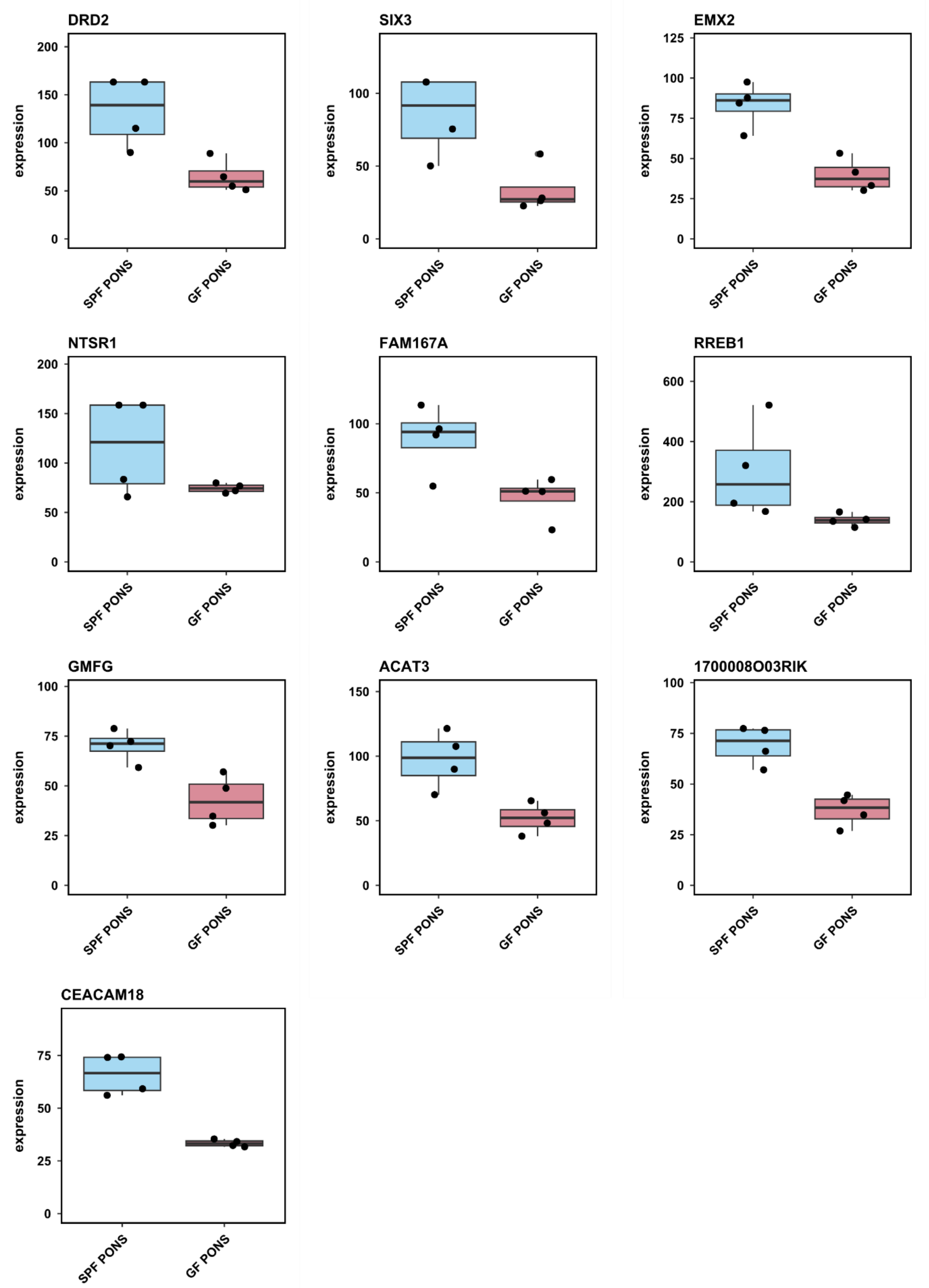

**Supplementary Figure 6. Boxplots of top ten DEGs with significantly lower expression in the GF compared to the SPF pons.** Boxplots of gene expression values of GF pons n=4 (red) and SPF pons n=4 (blue) for the ten most highly expressed DEGs found in GF pons, (*p*. adj < 0.05, absolute log2fold > 0.5). Each dot is one sample, with sample groups shown on x-axis and gene expression values on the y-axis with mean (parallel line) and standard error (vertical to x-axis) for each sample group.

**Supplementary Figure 7**

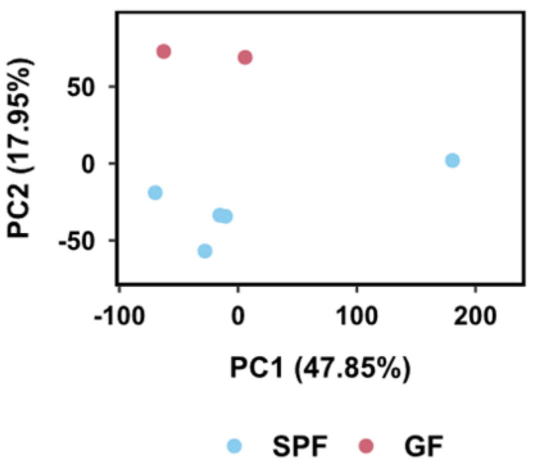


**Supplementary Figure 7. Sample clustering of DEGs between GF and SPF thalamus.** PCA plot of GF n=2 (red) and SPF n=5 (blue) thalamus samples, showing the first principal component (PC1) accounting for 47.85% of variance and the second principal component (PC2) capturing 17.95% of variance.

### Supplementary Figure 8

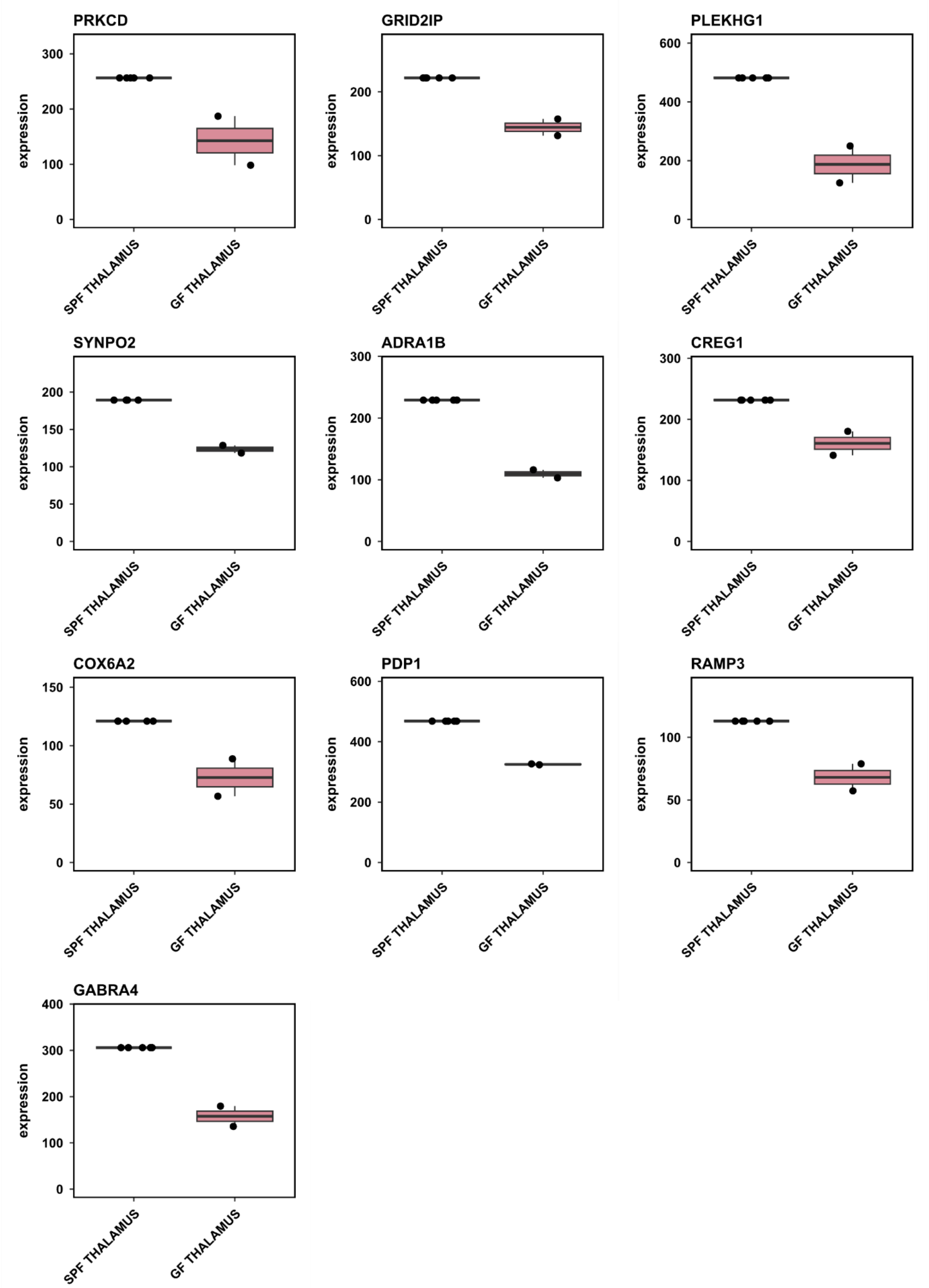

**Supplementary Figure 8. Boxplots of top ten DEGs with significantly lower expression in the GF compared to the SPF thalamus.** Boxplots of gene expression value of GF thalamus n=2 (red) and SPF thalamus n=5 (blue) for the ten most highly expressed genes found in GF thalamus by adjusted *p*-value, ($p < 0.05$). Each dot is one sample, with sample groups shown on x-axis and gene expression values on the y-axis with mean (parallel line) and standard error (vertical to x-axis) for each sample group.

**Supplementary Table 1**

| | Mice | | ROIs | |
|---|---|---|---|---|
| | **GF** | **SPF** | **GF** | **SPF** |
| Hippocampus | 3 | 5 | 7 | 17 |
| Thalamus | 2 | 5 | 2 | 7 |
| Pons | 4 | 4 | 5 | 6 |

**Supplementary Table 4. Summary table of mouse numbers for each brain region and number of ROIs selected across these samples.** In total 4 GF and 5 SPF mice were used in this study but for all mice it was not possible to identify ROIs due to sectioning artefacts. For example, 17 ROIs were selected for the SPF hippocampus across the 5 SPF mice.

**Supplementary Table 2**

| Gene | log2fold | p | p.adj |
|---|---|---|---|
| GPR68 | -1.11 | 8.016103e-09 | 7.730607e-06 |
| ADAMTS17 | -1.17 | 1.376545e-08 | 1.232696e-05 |
| LRRTM4 | -1.58 | 6.794341e-08 | 4.696261e-05 |
| PROX1 | -1.43 | 5.836582e-07 | 2.090664e-04 |
| MKX | -1.13 | 1.082724e-06 | 3.572135e-04 |
| CYP7B1 | -0.97 | 2.268044e-06 | 6.612668e-04 |
| PLEKHA2 | -1.10 | 2.427025e-06 | 6.915366e-04 |
| KCTD4 | -1.21 | 2.643806e-06 | 7.208062e-04 |
| GALNS | -1.06 | 2.778258e-06 | 7.298252e-04 |
| PTPRK | -1.02 | 3.610638e-06 | 8.230285e-04 |

**Supplementary Table 2. Top ten most significantly decreased DEGs in the GF compared to SPF hippocampus.** Ten most significantly decreased DEGs with cut off *p*. adj <0.05, absolute log2 fold > 0.5.

**Supplementary Table 3**

| Gene | log2fold | p | p.adj |
|---|---|---|---|
| DRD2 | -2.12 | 2.724298e-07 | 0.001446376 |
| SIX3 | -1.93 | 3.042012e-07 | 0.001446376 |
| EMX2 | -1.11 | 5.522510e-06 | 0.006059468 |
| NTSR1 | -1.68 | 3.254425e-05 | 0.029013201 |
| FAM167A | -1.11 | 1.139700e-04 | 0.067736182 |
| RREB1 | -1.11 | 1.314015e-04 | 0.074972447 |
| GMFG | -0.77 | 1.444158e-04 | 0.079228731 |
| ACAT3 | -0.92 | 1.684701e-04 | 0.086766276 |
| 1700008O03RIK | -0.94 | 2.261130e-04 | 0.099727515 |
| CEACAM18 | -1.01 | 2.955473e-04 | 1.000000000 |

**Supplementary Table 3. Top 10 DEGs significantly decreased in expression in the GF compared to SPF pons.** Ten most significant DEGs with cut off *p*. adj <0.05, absolute log2 fold >0.5.

**Supplementary Table 4**

| Gene | log2fold | p | p.adj |
|---|---|---|---|
| PRKCD | -4.06 | 2.895966e-21 | 3.380751e-17 |
| GRID2IP | -3.04 | 7.717696e-20 | 4.504819e-16 |
| PLEKHG1 | -2.94 | 7.536185e-19 | 2.932581e-15 |
| SYNPO2 | -3.01 | 3.020681e-16 | 8.815859e-13 |
| ADRA1B | -2.43 | 4.523080e-16 | 1.056049e-12 |
| CREG1 | -1.76 | 3.371579e-15 | 6.559969e-12 |
| COX6A2 | -3.00 | 1.079044e-14 | 1.799538e-11 |
| PDP1 | -2.32 | 2.492843e-14 | 3.637681e-11 |
| RAMP3 | -2.36 | 3.216601e-14 | 4.172289e-11 |
| GABRA4 | -2.64 | 8.405185e-14 | 9.812213e-11 |

**Supplementary Table 4. Top 10 DEGs significantly decreased in expression in the GF compared to SPF thalamus.** Significant DEGs with cut off *p*. adj <0.05, absolute log2 fold >0.5.
